# Caution when using network partners for target identification in drug discovery

**DOI:** 10.1101/2024.08.08.607024

**Authors:** Dandan Tan, Yiheng Chen, Yann Ilboudo, Kevin Y. H. Liang, Guillaume Butler-Laporte, J Brent Richards

## Abstract

Identifying novel, high-yield drug targets is challenging and often results in a high failure rate. However, recent data indicates that leveraging human genetic evidence to identify and validate these targets significantly increases the likelihood of success in drug development. Two recent papers from Open Targets claimed that around half of FDA-approved drugs had targets with direct human genetic evidence. By expanding target identification to include protein network partners—molecules in physical contact—the proportion of drug targets with genetic evidence support increased to two-thirds. However, the efficacy of using these network partners for target identification was not formally tested. To address this, we tested the approach on a list of robust positive control genes. We used the IntAct database to find molecular interacting proteins of genes identified by exome-wide association studies (ExWAS) and genome-wide association studies (GWAS) combined with a locus-to-gene mapping algorithm called the Effector Index (Ei). We assessed how accurately including interacting genes with the ExWAS and Effector Index selected genes identified positive controls, focusing on precision, sensitivity, and specificity. Our results indicated that although molecular interactions led to higher sensitivity in identifying positive control genes, their practical application is limited by low precision. Hence, expanding genetically identified targets to include network partners did not increase the chance of identifying drug targets, suggesting that such results should be interpreted with caution.

## Main Text

Using human genetics tools in drug discovery accelerates the identification of potential drug targets, but only a limited number of these targets have successfully translated into developed medicines^1-3^. The problems include the necessity for in vivo testing due to the complexities of biological systems and limitations in the number of available targets^4-6^. Network partners were defined as molecules in physical contact with each other at the same time^7,8^. While two recent papers from Open Targets suggested that more useful drug targets could be identified by expanding to network partners, it is not clear if this expansion would help to identify more viable drug development opportunities^9-12^. Specifically, the efficacy of including network partners to aid in target identification was not formally tested. Further, it is not known if the network partners of genetically identified targets are enriched for known disease-causal genes.

In this study, we utilized positive control genes (which were causes of Mendelian disease, or the targets of successfully developed medicines, described in Supplemental Methods) from a variety of sources that are known to influence disease risk, and tested whether adding IntAct-identified network partners can increase the ability and precision to find these positive controls, as depicted in Figure 1. The IntAct molecular interaction database is a curated repository of molecular interactions, sourced both from scientific literature and direct data submissions^7^. Briefly, to identify gene-trait pairs with support from human genetic evidence, we used ExWAS and GWAS combined with Effector Index^13,14^. The Effector Index algorithm is a computational tool designed to identify causal genes and provides a probability of causality for each gene at a GWAS locus, allowing for the ranking of each gene at such loci^14^. The outputs from exome sequencing and GWAS plus the Effector Index were then used to identify genetically-supported gene-trait pairs. We then tested whether these genetically-supported gene-trait pairs were enriched amongst the positive controls and if adding their network partners further improved identification of positive controls.

**Figure 1.**
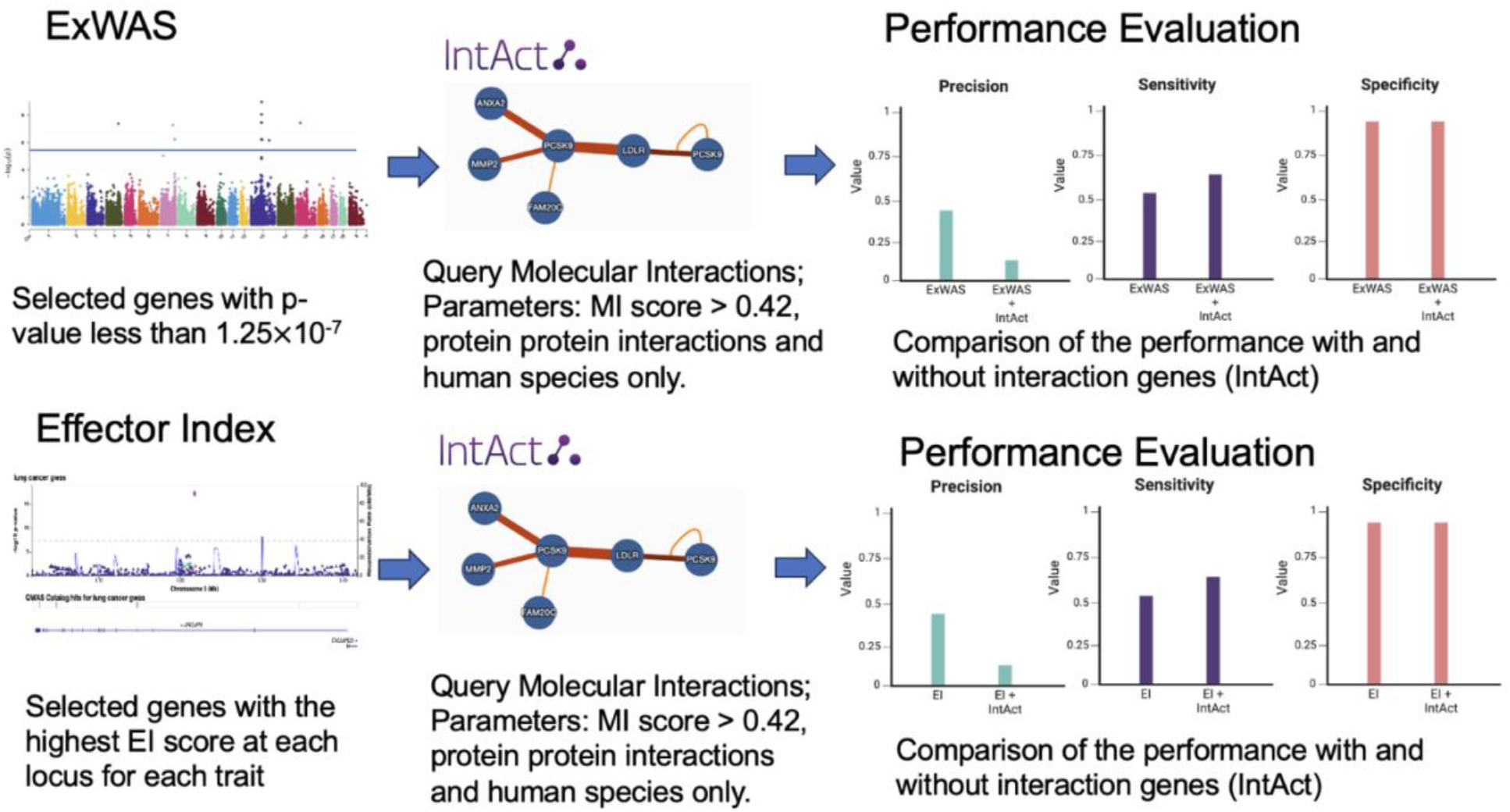
Workflow for combining molecular interactions via IntAct and evaluating the results. We utilized the ExWAS significant genes with a p-value less than 1.25×10^−7^ and identified the genes that interact with these significant genes using IntAct^13^. Then, we compared their performance to identify positive control genes by evaluating the sensitivity, specificity, and precision. Similarly, with the Effector Index (Ei) algorithm, we selected the genes with the highest Effector Index prediction score for each trait at each GWAS locus, found their molecular interacting genes using IntAct and evaluated their performance to identify positive control genes^14^.

To test whether network partners influence the precision to identify positive control genes, we began by focusing on the performance of ExWAS to identify these positive control genes.

We used the ExWAS burden test results from the UK biobank exome sequencing program with 12 continuous traits and 12 diseases. We selected the ExWAS significant genes as those with a p-value of less than 1.25 × 10^−7^ (0.05 / (20,000 genes * 4 variant masks * 5 different allele frequency inclusion thresholds))^13^. Next, we used the IntAct database to identify all molecular interactions of the resultant ExWAS significant genes using the same criteria as the Open Targets paper, which was a molecular interaction (MI) score greater than 0.42 in IntAct^9^. We compared the ability to identify positive control genes using only ExWAS significant genes with those derived from ExWAS significant genes and their network partners. Per variant mask, we found approximately 40 more positive control genes through network partners (Figure 2). However, this also resulted in a ten-fold increase in the number of interacting proteins whose genes were not positive control genes (Figure 2). We then assessed the precision, sensitivity, and specificity of ExWAS significant genes to identify positive control genes both including and excluding genes that are molecular interactors. While the sensitivity to identify positive control genes increased by 5%, there was a six-fold decrease in precision when including genes that were network partners compared to the ExWAS significant genes alone (Figure 3). Even when varying the MI threshold, the performance of inclusion of IntAct-identified genes remained poor (Table S1).

**Figure 2.**
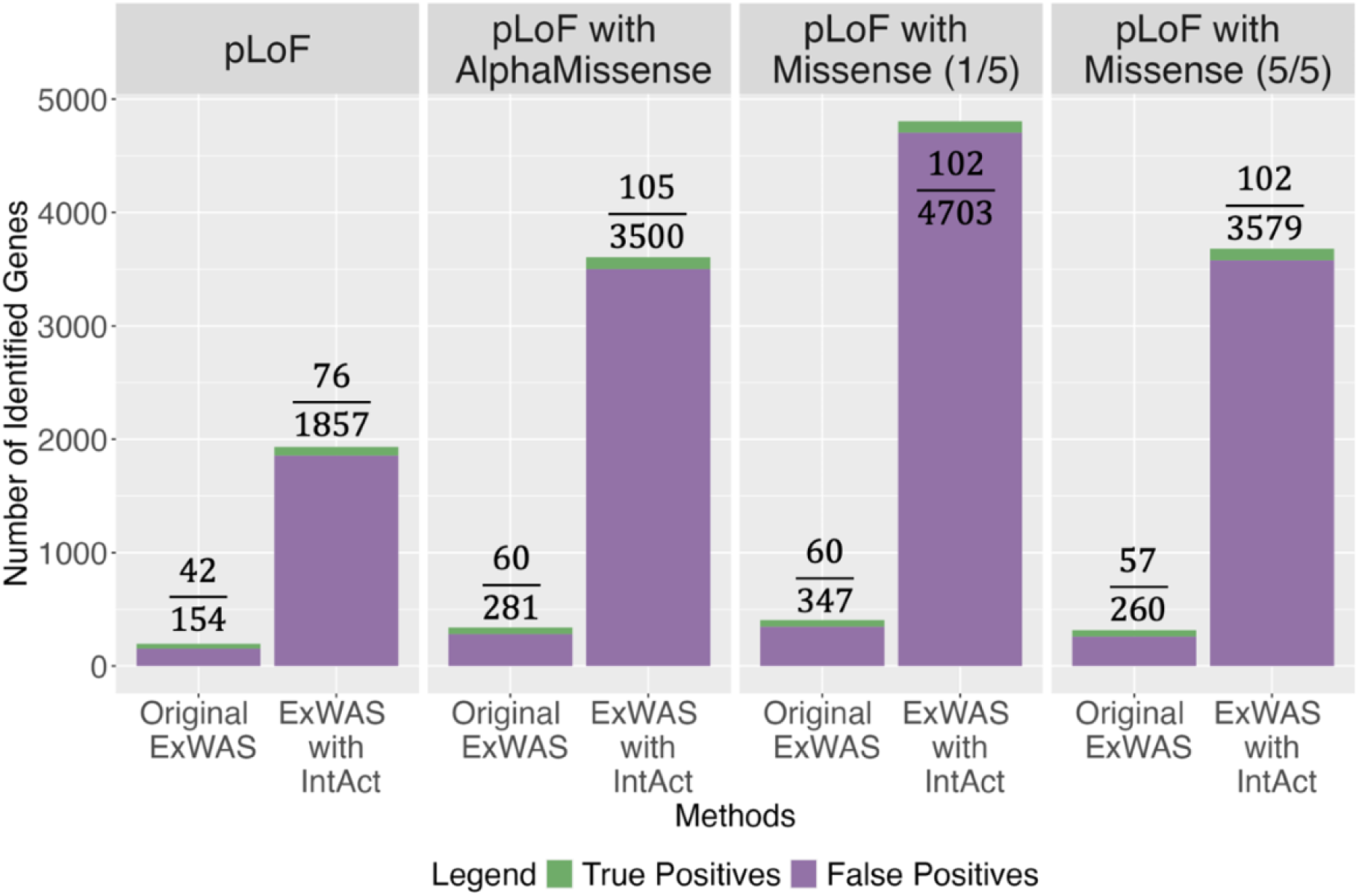
Identification of positive control genes across four ExWAS datasets. True positives (green) represented the ExWAS significant trait gene pairs, or their interactors also present in the positive control gene set. False positives (purple) represented the ExWAS significant trait gene pairs or their interactors that are not positive control genes.

**Figure 3.**
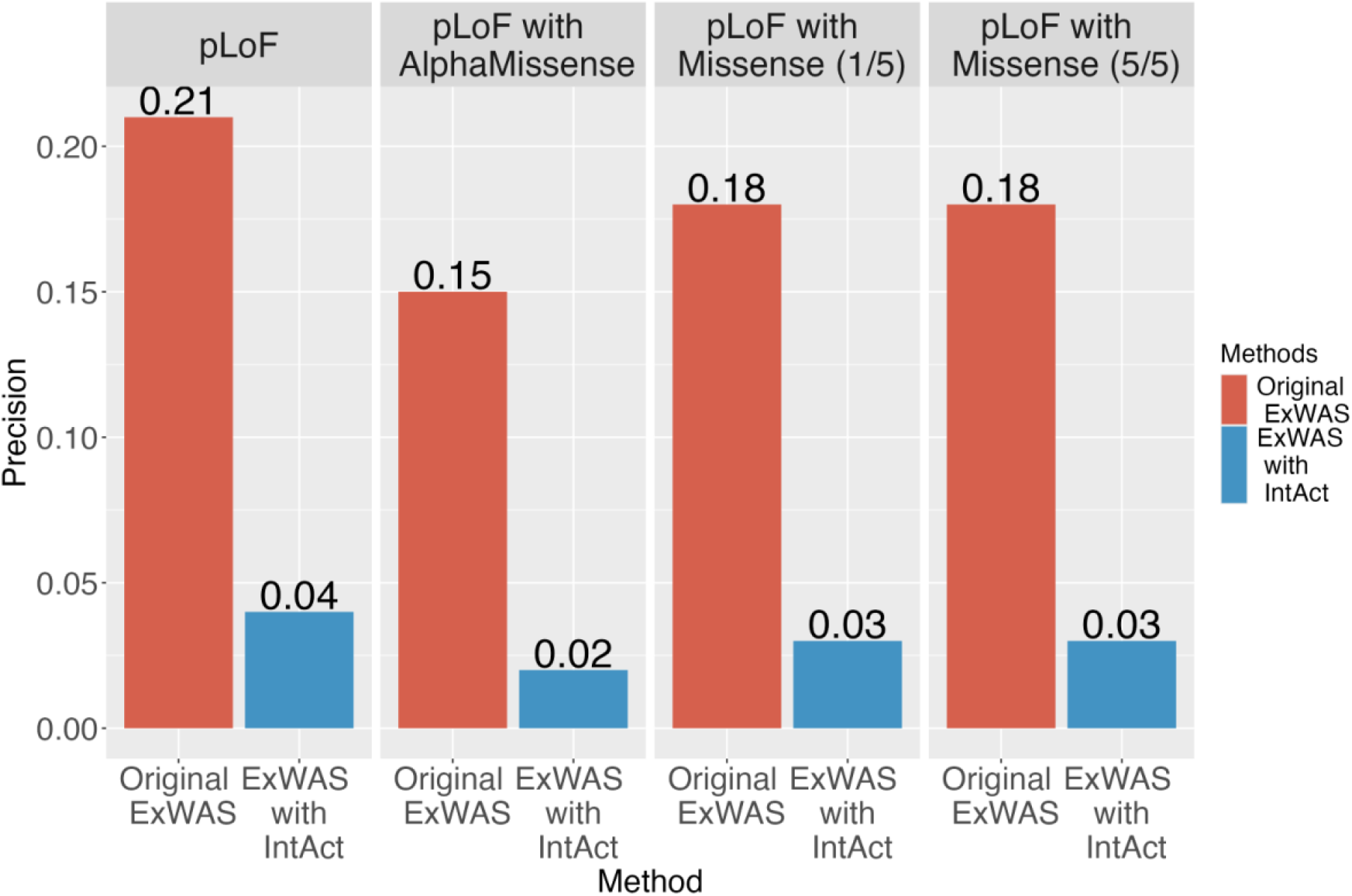
Precision comparison for positive control gene identification both including and excluding molecular interaction genes of ExWAS significant genes. The bar plot compares the precision of ExWAS significant genes (blue) with ExWAS significant genes and their interacting genes (red). ExWAS significant genes with their interacting genes exhibited lower precision to identify positive control genes.

We next studied the effect of adding network partners using the molecular interactions of genes identified from GWAS using the Effector Index prediction^14^. The Effector Index algorithm, trained on GWAS for 12 common diseases and quantitative traits, assigns scores ranging from 0 to 1 to all genes at a GWAS locus^14^. These scores aim to prioritize likely disease-causal genes at GWAS loci. A higher Effector Index score indicates a greater probability of a gene being a disease-causal gene. We selected the gene with the highest score for each trait at each locus. We only kept loci that contained at least one positive control gene since the absence of known targets would preclude our ability to evaluate the Effector Index’s performance. If not directly causal, we hypothesized that the interacting genes of those genes with the highest Effector Index score might be^15^. Using the IntAct database, we identified molecularly interacting genes at each GWAS locus with the highest Effector Index score, again using an MI score of 0.42 to identify interacting genes, again restricting our search to genes from humans^9^. We used the same positive control genes that were used in Effector Index study (Supplemental Methods)^14^. While the inclusion of genes with molecular interactions identified 47 additional positive control genes, it also included 1,370 additional genes that were not positive control genes (Figure 4). Consequently, the precision of Effector Index plus interacting protein predictions decreased by six-fold compared to the original Effector Index prediction algorithm (Figure 5). The sensitivity of Effector Index prediction combined with their molecular interactions in identifying positive control genes increased by 14%, and the specificity remained high. The poor performance in precision was because most interacting genes were not positive controls. Again, varying the MI threshold did not alter our results (Table S2).

**Figure 4.**
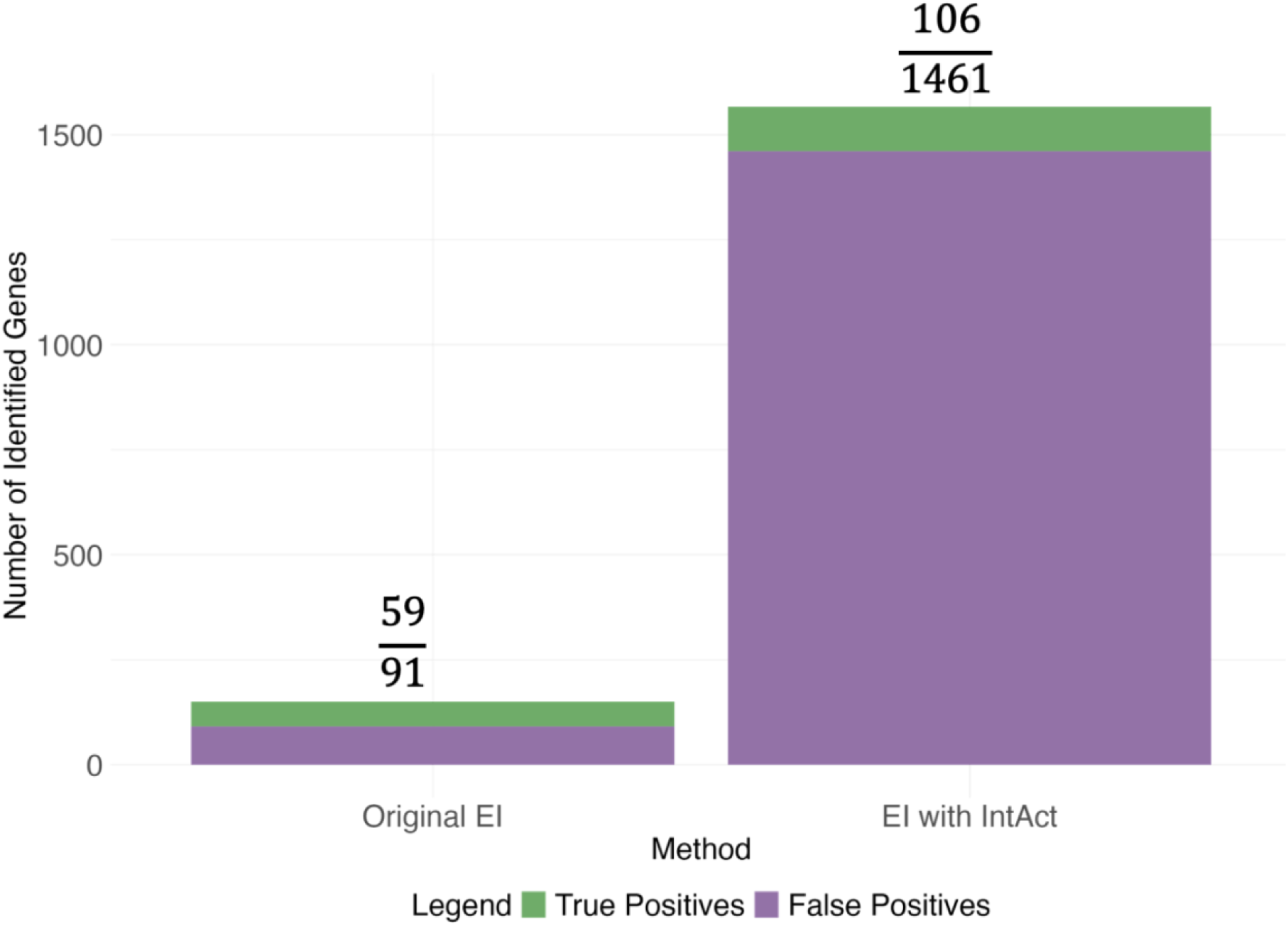
Identification of positive control genes by the Effector Index algorithm. This bar plot illustrates the performance of the Effector Index (Ei) prediction algorithm to identify positive control genes, comparing results with and without network partners. True positives are indicated in green, representing trait-gene pairs identified were positive control genes. False positives are represented in purple, representing trait gene pairs identified by the algorithm that were not positive control genes.

**Figure 5.**
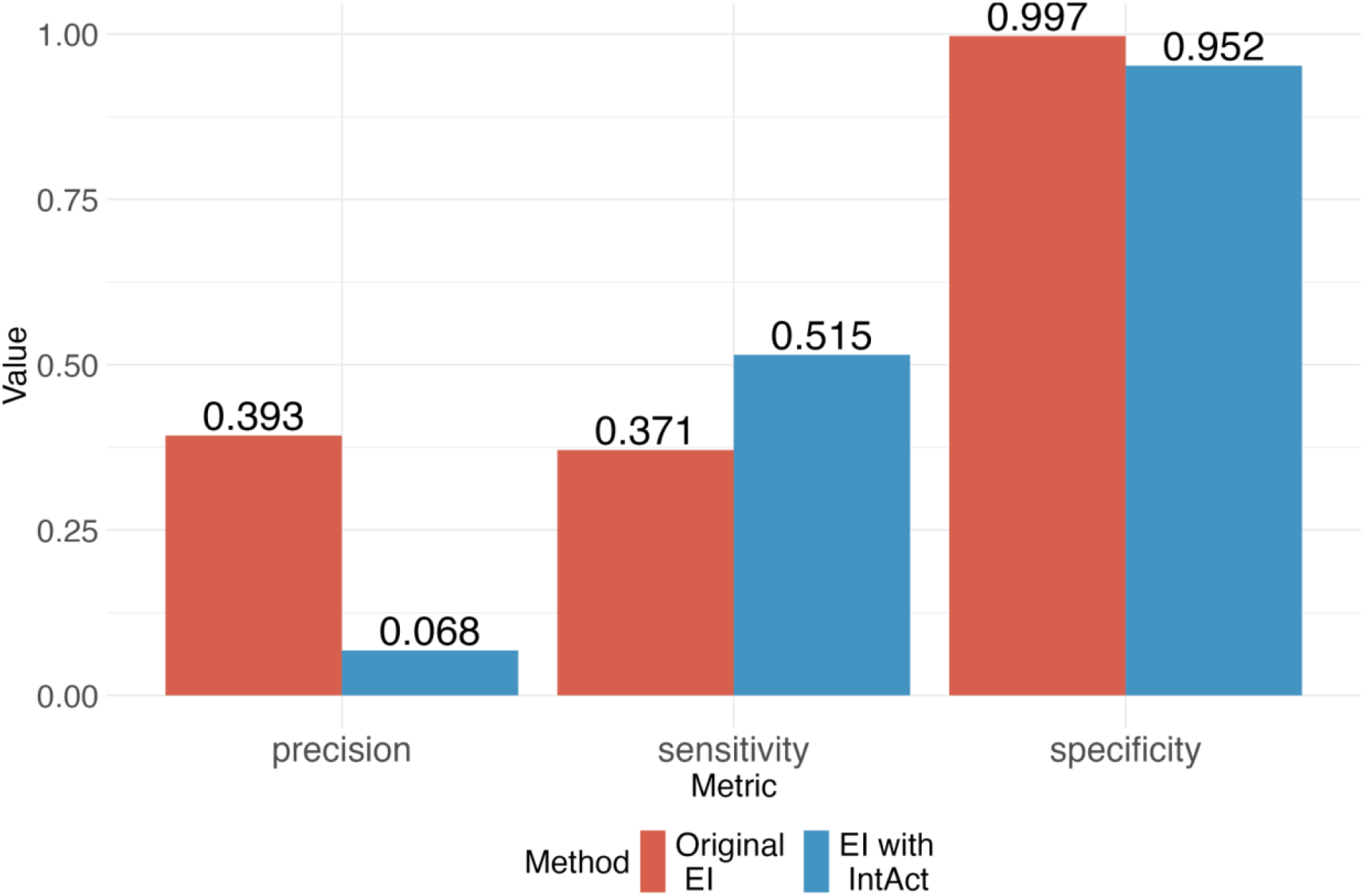
Performance comparison of positive control genes identification both including and excluding molecular interaction genes of Effector Index algorithm. The bar plot compared the precision, sensitivity, and specificity of the Effector Index prediction algorithm to identify positive control genes with and without network partners. Incorporating network partners resulted in lower precision and specificity but higher sensitivity compared to the baseline results.

Taken together, while the Open Target studies have highlighted the potential of molecular interactions to identify high-yield drug targets, their practical application is hindered by low precision. This makes it challenging to employ molecular interactors in real-world scenarios like the identification of new drug targets. Among the interactions we identified, only a small fraction were positive controls; discerning these valuable few from the majority identified proved difficult resulting in a low precision to identify positive controls. In conclusion, expanding genetically identified targets by including their network partners does not appear to increase the ability to identify positive control genes and such results should be interpreted with caution.

## Data and Code Availability

The Open Target data was downloaded from (February 2^nd^, 2024): https://platform.opentargets.org/downloads (version 2021-12)

The code and data necessary to reproduce the figures can be found here: https://github.com/richardslab/IntAct_Report.git

**Data for ExWAS** (https://github.com/richardslab/IntAct_Report/tree/main/ExWAS_Data): Positive control gene list of ExWAS

Significant genes list of pLoF dataset

Significant genes list of pLoF with AlphaMissense

Significant genes list of pLoF with Missense (5/5) dataset

Significant genes list of pLoF with Missense (1/5) dataset

**Data for Effector Index Prediction**

(https://github.com/richardslab/IntAct_Report/tree/main/EI_Data): Positive control gene list of Effector Index Prediction Effector Index prediction results for all traits

## Acknowledgments

The Richards research group is supported by the Canadian Institutes of Health Research (CIHR: 365825, 409511, 100558, 169303), the McGill Interdisciplinary Initiative in Infection and Immunity (MI4), the Lady Davis Institute of the Jewish General Hospital, the Jewish General Hospital Foundation, the Canadian Foundation for Innovation, the NIH Foundation, Cancer Research UK, Genome Québec, the Public Health Agency of Canada, McGill University, Cancer Research UK, and the Fonds de Recherche Québec Santé (FRQS). J.B.R. is supported by an FRQS Mérite Clinical Research Scholarship. Support from Calcul Québec and Compute Canada is acknowledged. TwinsUK is funded by the Welcome Trust, Medical Research Council, European Union, the National Institute for Health Research (NIHR)-funded BioResource, Clinical Research Facility and Biomedical Research Centre based at Guy’s and St Thomas’ NHS Foundation Trust in partnership with King’s College London. D.T is supported by Quantitative Life Sciences of McGill University. GBL is supported by CIHR and FRQS salary support grants.

## Author Contribution Statement

DT - Writing initial draft. DT, YC, KYHL, GBL, JBR – Methodology. DT, YC - Data Analysis. DT, YC, YI, GBL, JBR - Writing review and editing draft. JBR – Supervision. All authors commented/revised the manuscript and agreed to its final submitted version.

## Reference

1. Somda, D., Wilson Kpordze, S., Jerpkorir, M., Chantelle Mahora, M., Wanjiru Ndungu, J., Wambui Kamau, S., Arthur, V., and Elbasyouni, A. (2023). The Role of Bioinformatics in Drug Discovery: A Comprehensive Overview. In Drug Metabolism and Pharmacokinetics (IntechOpen). 10.5772/intechopen.113712.

2. Saharan, V.A. ed. (2022). Computer Aided Pharmaceutics and Drug Delivery: An Application Guide for Students and Researchers of Pharmaceutical Sciences (Springer Nature Singapore) 10.1007/978-981-16-5180-9.

3. Walker, M.J.A., Barrett, T., and Guppy, L.J. (2004). Functional pharmacology: the drug discovery bottleneck? Drug Discovery Today: TARGETS 3, 208–215. 10.1016/S1741-8372(04)02449-1.

4. Hormozdiari, F., Kichaev, G., Yang, W.-Y., Pasaniuc, B., and Eskin, E. (2015). Identification of causal genes for complex traits. Bioinformatics 31, i206–i213. 10.1093/bioinformatics/btv240.

5. Schork, N.J. (1997). Genetics of Complex Disease. Am J Respir Crit Care Med 156, S103–S109. 10.1164/ajrccm.156.4.12-tac-5.

6. MacNamara, A., Nakic, N., Amin Al Olama, A., Guo, C., Sieber, K.B., Hurle, M.R., and Gutteridge, A. (2020). Network and pathway expansion of genetic disease associations identifies successful drug targets. Sci Rep 10, 20970. 10.1038/s41598-020-77847-9.

7. del Toro, N., Shrivastava, A., Ragueneau, E., Meldal, B., Combe, C., Barrera, E., Perfetto, L., How, K., Ratan, P., Shirodkar, G., et al. (2022). The IntAct database: efficient access to fine-grained molecular interaction data. Nucleic Acids Research 50, D648–D653. 10.1093/nar/gkab1006.

8. Barabási, A.-L., Gulbahce, N., and Loscalzo, J. (2011). Network medicine: a network-based approach to human disease. Nat Rev Genet 12, 56–68. 10.1038/nrg2918.

9. Ochoa, D., Karim, M., Ghoussaini, M., Hulcoop, D.G., McDonagh, E.M., and Dunham, I. (2022). Human genetics evidence supports two-thirds of the 2021 FDA-approved drugs. Nat Rev Drug Discov 21, 551–551. 10.1038/d41573-022-00120-3.

10. Rusina, P.V., Falaguera, M.J., Romero, J.M.R., McDonagh, E.M., Dunham, I., and Ochoa, D. (2023). Genetic support for FDA-approved drugs over the past decade. Nat Rev Drug Discov 22, 864–864. 10.1038/d41573-023-00158-x.

11. Loscalzo, J. (2023). Molecular interaction networks and drug development: Novel approach to drug target identification and drug repositioning. The FASEB Journal 37, e22660. 10.1096/fj.202201683R.

12. Cheng, F., Kovács, I.A., and Barabási, A.-L. (2019). Network-based prediction of drug combinations. Nat Commun 10, 1197. 10.1038/s41467-019-09186-x.

13. Chen, Y., Butler-Laporte, G., Liang, K.Y.H., Ilboudo, Y., Yasmeen, S., Sasako, T., Langenberg, C., Greenwood, C.M.T., and Richards, J.B. (2024). The performance of AlphaMissense to identify genes causing disease (Genetic and Genomic Medicine) 10.1101/2024.03.05.24303647.

14. Forgetta, V., Jiang, L., Vulpescu, N.A., Hogan, M.S., Chen, S., Morris, J.A., Grinek, S., Benner, C., Jang, D.-K., Hoang, Q., et al. (2022). An effector index to predict target genes at GWAS loci. Hum Genet 141, 1431–1447. 10.1007/s00439-022-02434-z.

15. Mountjoy, E., Schmidt, E.M., Carmona, M., Schwartzentruber, J., Peat, G., Miranda, A., Fumis, L., Hayhurst, J., Buniello, A., Karim, M.A., et al. (2021). An open approach to systematically prioritize causal variants and genes at all published human GWAS trait-associated loci. Nat Genet 53, 1527–1533. 10.1038/s41588-021-00945-5.

