## Supplemental file for "Caution when using network partners for target identification in drug discovery"

### Supplemental information

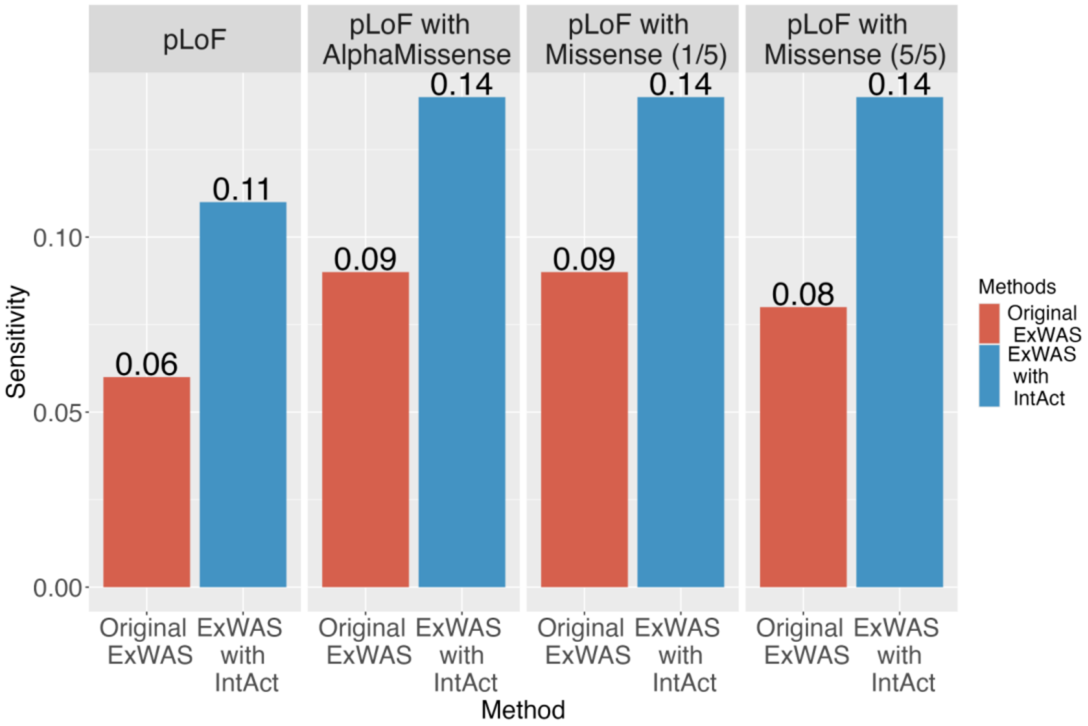


Figure S1. Sensitivity comparison for positive control gene identification both including and excluding molecular interaction genes of ExWAS significant genes.

The bar plot compares the sensitivity of ExWAS significant genes with and without their interacting genes. ExWAS significant genes with their interacting genes had a higher sensitivity to identify positive control genes.

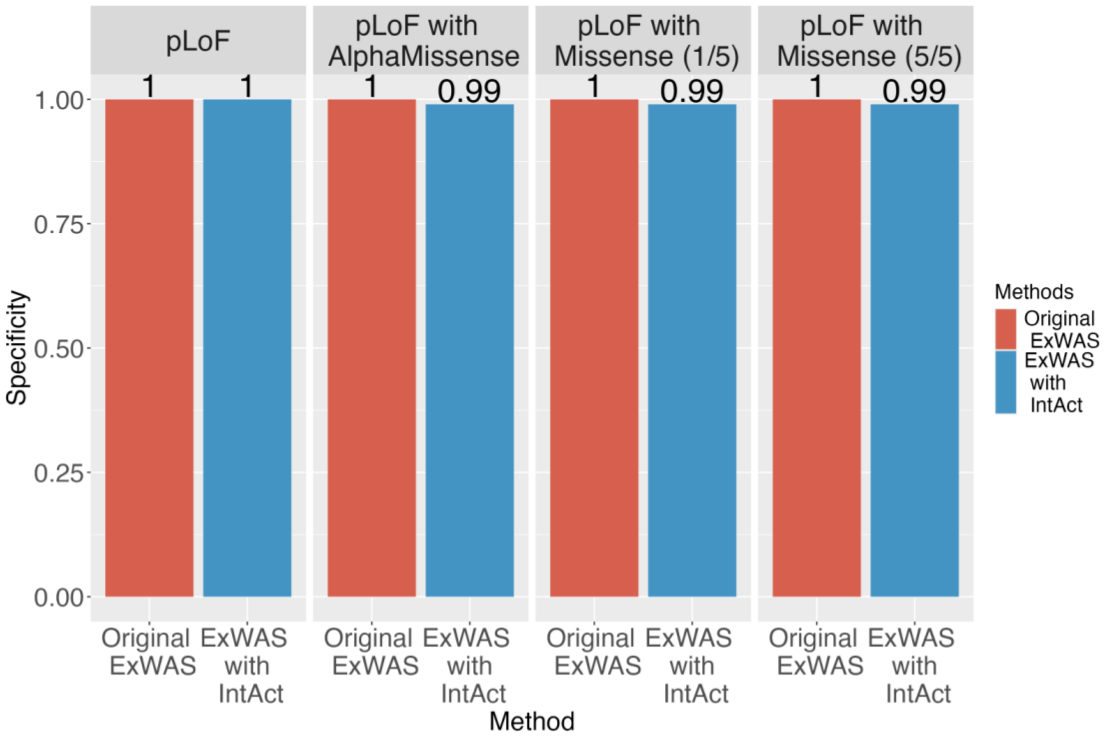


Figure S2. Specificity comparison for positive control gene identification both including and excluding molecular interaction genes of ExWAS significant genes.

The bar plot compares the specificity of ExWAS significant genes with and without their interacting. ExWAS significant genes with their interacting genes had similar specificity across all four datasets. The values are rounded to 2 significant figures.

Table S1. Performance metrics at varying MI scores when including interactors of four ExWAS methods.

A. pLoF dataset

| MI score | 0 | 0.1 | 0.2 | 0.3 | 0.4 | 0.5 | 0.6 | 0.7 | 0.8 | 0.9 | 1 |
|---|---|---|---|---|---|---|---|---|---|---|---|
| Precision | 0.015 | 0.015 | 0.015 | 0.017 | 0.039 | 0.041 | 0.129 | 0.162 | 0.200 | 0.216 | 0.214 |
| Sensitivity | 0.136 | 0.136 | 0.136 | 0.135 | 0.106 | 0.097 | 0.083 | 0.076 | 0.067 | 0.066 | 0.062 |
| Specificity | 0.983 | 0.983 | 0.983 | 0.985 | 0.995 | 0.996 | 0.998 | 0.999 | 0.999 | 0.999 | 0.999 |

B. pLoF with AlphaMissense

| MI score | 0 | 0.1 | 0.2 | 0.3 | 0.4 | 0.5 | 0.6 | 0.7 | 0.8 | 0.9 | 1 |
|---|---|---|---|---|---|---|---|---|---|---|---|
| Precision | 0.014 | 0.014 | 0.014 | 0.015 | 0.029 | 0.030 | 0.106 | 0.126 | 0.165 | 0.176 | 0.176 |
| Sensitivity | 0.181 | 0.181 | 0.181 | 0.178 | 0.141 | 0.127 | 0.110 | 0.095 | 0.091 | 0.090 | 0.086 |
| Specificity | 0.975 | 0.975 | 0.975 | 0.977 | 0.991 | 0.992 | 0.998 | 0.999 | 0.999 | 0.999 | 0.999 |

C. pLoF with Missense (1/5)

| MI score | 0 | 0.1 | 0.2 | 0.3 | 0.4 | 0.5 | 0.6 | 0.7 | 0.8 | 0.9 | 1 |
|---|---|---|---|---|---|---|---|---|---|---|---|
| Precision | 0.011 | 0.011 | 0.011 | 0.012 | 0.028 | 0.030 | 0.106 | 0.128 | 0.159 | 0.179 | 0.180 |
| Sensitivity | 0.171 | 0.171 | 0.171 | 0.171 | 0.138 | 0.129 | 0.109 | 0.096 | 0.087 | 0.086 | 0.082 |
| Specificity | 0.971 | 0.971 | 0.971 | 0.972 | 0.991 | 0.992 | 0.998 | 0.999 | 0.999 | 0.999 | 0.999 |

D. pLoF with Missense (5/5)

| MI score | 0 | 0.1 | 0.2 | 0.3 | 0.4 | 0.5 | 0.6 | 0.7 | 0.8 | 0.9 | 1 |
|---|---|---|---|---|---|---|---|---|---|---|---|
| Precision | 0.009 | 0.009 | 0.009 | 0.010 | 0.021 | 0.022 | 0.079 | 0.091 | 0.134 | 0.147 | 0.147 |
| Sensitivity | 0.173 | 0.173 | 0.173 | 0.169 | 0.138 | 0.126 | 0.111 | 0.095 | 0.091 | 0.090 | 0.086 |
| Specificity | 0.966 | 0.966 | 0.966 | 0.968 | 0.988 | 0.990 | 0.998 | 0.998 | 0.999 | 0.999 | 0.999 |

Table S2. Performance metrics at varying MI scores for including interactors of the Effector Index algorithm.

| MI score | 0 | 0.1 | 0.2 | 0.3 | 0.4 | 0.5 | 0.6 | 0.7 | 0.8 | 0.9 | 1 |
|---|---|---|---|---|---|---|---|---|---|---|---|
| Precision | 0.024 | 0.024 | 0.024 | 0.026 | 0.067 | 0.071 | 0.222 | 0.286 | 0.349 | 0.375 | 0.393 |
| Sensitivity | 0.590 | 0.590 | 0.590 | 0.581 | 0.515 | 0.482 | 0.451 | 0.422 | 0.394 | 0.387 | 0.371 |
| Specificity | 0.831 | 0.831 | 0.831 | 0.845 | 0.951 | 0.959 | 0.990 | 0.994 | 0.996 | 0.996 | 0.997 |

### Supplemental Methods

**ExWAS datasets**: ExWAS focuses on identifying coding variants associated with specific traits or diseases. The exome comprises all exons in a genome, representing the DNA sequences that are translated into proteins. Despite being a small portion of the human genome, the exome harbors a disproportionate number of known disease-associated variants[1]. We used all four ExWAS results obtained from Chen et al. (2024). His method used whole-exome sequencing data from the UK Biobank to compare four gene-based association analysis methods, incorporating deleterious genetic variants: pLoF variants only (pLoF), the combination of pLoF with AlphaMissense pathogenic variants (pLoF with AlphaMissense), pLoF with five commonly utilized annotation methods that predict deleterious variants (pLoF with Missense (5/5)) and pLoF with only variants predicted to be deleterious by all five methods (pLoF with Missense (1/5)). We chose ExWAS significant genes with a p-value less than $1.25 \times 10^{-7}$ (0.05 / (20,000 genes * 4 variant masks * 5 different allele frequency inclusion thresholds))[2].

**Positive Control Genes for ExWAS:** The performance evaluation relied on a positive control gene list containing 900 disease gene pairs, known to cause Mendelian diseases or be targets of successful drug development[2]. This positive control genes list was extracted from studies that used these genes to train algorithms to prioritize disease-causal or drug-targeting genes from GWAS signals[3,4]. They created their positive control gene lists by integrating genetic evidence, drug-target-indication associations, and expert manual curation from board-certified physicians and specialists. The positive control gene list also included Mendelian diseases genes from the MendelVar database. By combining these diverse sources of information, a list of positive control genes was constructed and used for evaluating the performance of ExWAS. The full list of positive control genes for each trait and disease can be found on GitHub (https://github.com/richardslab/IntAct_Report).

**Effector Index**: The Effector Index algorithm is a computational tool designed to identify causal genes at GWAS loci[3]. This algorithm employs fine-mapping of GWAS summary statistics, followed by the annotation of single nucleotide variants (SNVs) to identify potential causal variants and their association with genes within or near the GWAS loci. Subsequently, gene- and locus-level features are generated by integrating information on the presence of specific SNVs within genes, their functional implications, and locus-specific characteristics. Feature weights within the models were determined through leave-one-out analysis, enabling the assessment of model robustness and generalizability. Genomic features with higher Effector Index scores are considered more likely to be causal. We chose the genes with highest Effector Index scores at each locus for each trait.

**Positive Control Genes for Effector Index:** We used the same set of positive control genes used in the Effector Index study[3]. The Effector Index study utilized two approaches to define this list. Firstly, clinician scientists manually curated the Human Disease Ontology database to identify relevant ontological terms, and then utilized associated OMIM (Online Mendelian Inheritance in Man) linkage information to compile a list of genes associated with these diseases. Secondly, clinician scientists identified drug targets for medications with known mechanisms of action. Additionally, they incorporated data from the National Institutes of Health Type 2 Diabetes Accelerated Medicines Program, a collaborative effort between industry and academia aimed at identifying causal genes for Type 2 Diabetes. To ensure the reliability of the positive control genes list, genes labeled as "causal" solely based on evidence from GWAS were excluded, as their inclusion could introduce bias in the evaluation of Effector Index's performance. The complete list of positive control genes for the Effector Index of each trait is provided on GitHub (https://github.com/richardslab/IntAct_Report).

**IntAct Database**: The IntAct molecular interaction database serves as a curated repository of molecular interactions, drawing data from scientific literature and direct submissions[4]. These interactions are annotated, considering factors such as the detection method, interaction type, and the number of publications reporting a specific interaction. This information is consolidated into a Molecular Interaction (MI) score, reflecting the confidence level in the existence of each interaction within the dataset. The MI score provides a standardized measure of reliability for molecular interactions. In our study, we filtered interactions to retain only those with an MI score greater than 0.42, as specified in the Open Targets paper[5]. This threshold is thought to ensure a sufficient level of confidence in the interactions selected for analysis. Additionally, to maintain consistency with the genes listed in the ExWAS and Effector Index prediction datasets, we restricted the interactions to those involving human genes exclusively.

**Performance Evaluation**: We defined True Positive (TP), False Positive (FP), True Negative (TN), and False Negative (FN) as follows. For ExWAS datasets, TPs were genes with significant p-values or their interactors present in the positive control gene list; FPs were genes with significant p-values or their interactors not present in the positive control gene list; TNs were genes with nonsignificant p-values and not present in the positive control gene list; FNs were genes with nonsignificant p-values but present in the positive control gene list. Similar definitions were applied for the Effector Index algorithm. TPs were genes with the highest Effector Index scores at each locus for each trait or their interactors present in the positive control gene list; FPs were genes with the highest Effector Index scores at each locus for each trait or their interactors not present in the positive control gene list; TNs were genes with low Effector Index scores (< 0.46) and not present in the positive control gene list[3]; FNs were genes with low Effector Index scores (< 0.46) but present in the positive control gene list. We then calculated precision (TP / (TP + FP)), sensitivity (TP / (TP + FN)), and specificity (TN / (TN + FP)) to evaluate the performance of adding interacting genes from the IntAct database.

### Supplemental References


1. Zhang, L., Yu, L., Shu, X., Ding, J., Zhou, J., Zhong, C., Pan, B., Guo, W., Zhang, C., and Wang, B. (2023). Whole exome sequencing reveal 83 novel Mendelian disorders carrier P/LP variants in Chinese adult patients. J Hum Genet *68*, 737–743. https://doi.org/10.1038/s10038-023-01179-5.
2. Chen, Y., Butler-Laporte, G., Liang, K.Y.H., Ilboudo, Y., Yasmeen, S., Sasako, T., Langenberg, C., Greenwood, C.M.T., and Richards, J.B. (2024). The performance of AlphaMissense to identify genes causing disease (Genetic and Genomic Medicine) https://doi.org/10.1101/2024.03.05.24303647.
3. Forgetta, V., Jiang, L., Vulpescu, N.A., Hogan, M.S., Chen, S., Morris, J.A., Grinek, S., Benner, C., Jang, D.-K., Hoang, Q., et al. (2022). An effector index to predict target genes at GWAS loci. Hum Genet *141*, 1431–1447. https://doi.org/10.1007/s00439-022-02434-z.
4. del Toro, N., Shrivastava, A., Ragueneau, E., Meldal, B., Combe, C., Barrera, E., Perfetto, L., How, K., Ratan, P., Shirodkar, G., et al. (2022). The IntAct database: efficient access to fine-grained molecular interaction data. Nucleic Acids Research 50, D648–D653. https://doi.org/10.1093/nar/gkab1006.
5. Ochoa, D., Karim, M., Ghoussaini, M., Hulcoop, D.G., McDonagh, E.M., and Dunham, I. (2022). Human genetics evidence supports two-thirds of the 2021 FDA-approved drugs. Nat Rev Drug Discov *21*, 551–551. https://doi.org/10.1038/d41573-022-00120-3.
